# Hypoxia regulates endogenous double-stranded RNA production via reduced mitochondrial DNA transcription

**DOI:** 10.1101/2020.07.31.230300

**Authors:** Esther Arnaiz, Ana Miar, Antonio Gregorio Dias, Naveen Prasad, Ulrike Schulze, Dominic Waithe, Jan Rehwinkel, Adrian L. Harris

## Abstract

Hypoxia is a common phenomenon in solid tumours strongly linked to the hallmarks of cancer. Hypoxia promotes local immunosuppression and downregulates type I interferon (IFN) expression and signalling, which contribute to the success of many cancer therapies. Double-stranded RNA (dsRNA), transiently generated during mitochondrial transcription, endogenously activates the type I IFN pathway. We report the effects of hypoxia on the generation of mitochondrial dsRNA (mtdsRNA) in breast cancer. We found a significant decrease in dsRNA production in different cell lines under hypoxia. This was HIF1α/2α-independent. mtdsRNA was responsible for induction of type I IFN and significantly decreased after hypoxia. Mitochondrially encoded gene expression was downregulated and mtdsRNA bound by the dsRNA-specific J2 antibody was decreased during hypoxia. These findings reaveal a mechanism of hypoxia-induced immunosuppression that could be targeted by hypoxia-activated therapies.

## INTRODUCTION

Type I interferons (IFNs) include 13 IFNα subtypes, IFNβ, IFNε, IFNκ and IFNω, and type II and type III IFNs include IFNγ and IFNλ1-4, respectively. All IFNs are involved in the innate immune response against pathogenic infection. Type I and III IFNs are induced when specific microbial products, known as pathogen-associated molecular patterns (PAMPs), are detected pattern-recognition receptors (PRRs)^1,2^. PRRs include the Toll-like receptors (TLRs), some of which are specialised to survey the endosomal compartment for nucleic acids. In the cytosol, retinoic acid-inducible gene I (RIG-I) and melanoma differentiation-associated gene 5 (MDA5) detect unusual RNA molecules, while cyclic GMP-AMP (cGAMP) synthase (cGAS) is the major cytosolic double-strand DNA (dsDNA) sensor and activates stimulator of interferon genes (STING).

Recognition of viral RNA by RIG-I and MDA5 induces protein conformational changes, which allow interaction with the shared adaptor mitochondrial antiviral-signaling protein (MAVS) that then triggers phosphorylation of interferon-regulatory factor 3 (IRF3) and IRF7. These transcription factors induce the expression of type I and III IFNs, chemokines, inflammatory cytokines and other genes^3^. All type I IFNs bind a common receptor formed by IFNAR1 and IFNAR2, which through tyrosine kinase 2 (TKY2) and Janus kinase 1 (JAK1) recruits and phosphorylates signal transducer and activator of transcription (STAT) proteins^4^. The canonical IFNAR signalling cascade involves STAT1 and STAT2, which form a ternary complex called interferon-stimulated gene factor 3 (ISGF3) with interferon-regulatory factor 9 (IRF9). ISGF3 translocates to the nucleus where it activates the transcription of IFN-stimulated genes (ISGs)^2^. Interestingly, type I IFNs can be produced in the absence of infectionand are involved in the success of many anticancer treatments such as radiotherapy, chemotherapy, immunotherapy and oncolytic viruses^5^, promoting direct (tumour cell inhibition) and indirect (antitumour immune response) effects^6^.

Hypoxia generates an immunosuppressive microenvironment within the tumour by impeding the homing of immune effector cells and blocking their activity^7^. Additionally, tumours contain more immunosuppressive cells, such as myeloid-derived suppressor cells (MDSCs), tumour-associated macrophages (TAMs) and T-regulatory (Treg) cells, in hypoxic regions^7,8^.

Lactate, generated during the metabolic switch to glycolysis under hypoxia, acts as a ‘signalling molecule’ and attenuates the cytotoxic activity of CTLs^9^ and NK cells^10^, helps recruit MDSCs to the tumour^10^, and inhibits type I IFN induction via MAVS^11^.

Mitochondrial DNA (mtDNA) is a closed-circular, dsDNA molecule of about 16.6 kb^12^. Its complementary DNA strands are called H (heavy) and L (light). The H strand encodes 28 genes: 2 rRNAs, 14 tRNAs and 12 polypeptides, whereas the L strand only contains 9 genes encoding for 8 tRNAs and a single polypeptide. Mitochondria generate a number of damage-associated molecular patterns (DAMPs) including ATP, succinate, cardiolipin, N-formylpeptides, mitochondrial transcription factor A (TFAM), cytochrome-c, mtDNA and mitochondrial RNA (mtRNA)^13^. Extracellular mtDNA binds to TRL9^14^ whereas cytosolic mtDNA is recognised by inflammasomes^15^ and cGAS^16^. More recently, mtRNA was described to be a potent DAMP via recognition of a specific segment of the mitochondrial single-strand rRNA by TLR8^17^. In addition, dsRNA originating from convergent mtDNA transcription triggers an MDA5-dependent type I IFN response when released to the cytoplasm^18^.

Previously, we showed that the type I IFN responses induced by exogenous dsRNA are downregulated under hypoxia in cell lines from different solid tumours via transcriptional repression after changes in chromatin conformation^19^. Here, we investigated the role of hypoxia in the regulation of endogenous dsRNA formation and function. We have shown that hypoxia descreases the formation of mtdsRNA probably by reducing the mitochondrial transcription rate, and consequently lower endogenous activation of the type I IFN pathway. This effect is HIF1α/2α independent and occurs in different cancer cell lines as well as non-transformed cell lines. Moreover, different tissues have different immunostimulatory potential.

## RESULTS

### Hypoxia prevents the accumulation of immunostimulatory RNAs

Given the immuno-suppressive role of hypoxia, we tested whether cancer cell lines cultured in normoxia or hypoxia contain different amounts of immunostimulatory RNA. We extracted total RNA from the breast cancer cell line MCF7, cultured for 48 hours in normoxia or in 1% or 0.1% hypoxia We then transfected this RNA into an *IFNβ* promoter reporter cell line^20^. RNA from cells grown in normoxia induced expression of the reporter, indicative of the presence of immunostimlautory RNA (fig. 1a). To determine the sensor for this endogenous RNA, we tested the response in reporter cells lacking MDA5, RIG-I or MAVS. This analysis showed that total RNA from MCF7 cells induced an MDA5-MAVS-dependent response (fig 1a). Interestingly, RNA from hypoxic cells (hypoxic RNA) had a significantly reduced capacity to induce activation of the *IFNβ* promoter reporter compared with RNA from normoxic cells (normoxic RNA; fig. 1a). Moreover, much like the response to normoxic RNA, residual reporter induction after hypoxic RNA transfection was MDA5-MAVS-dependent and RIG-I-independent (fig. 1a).

**Figure 1.**
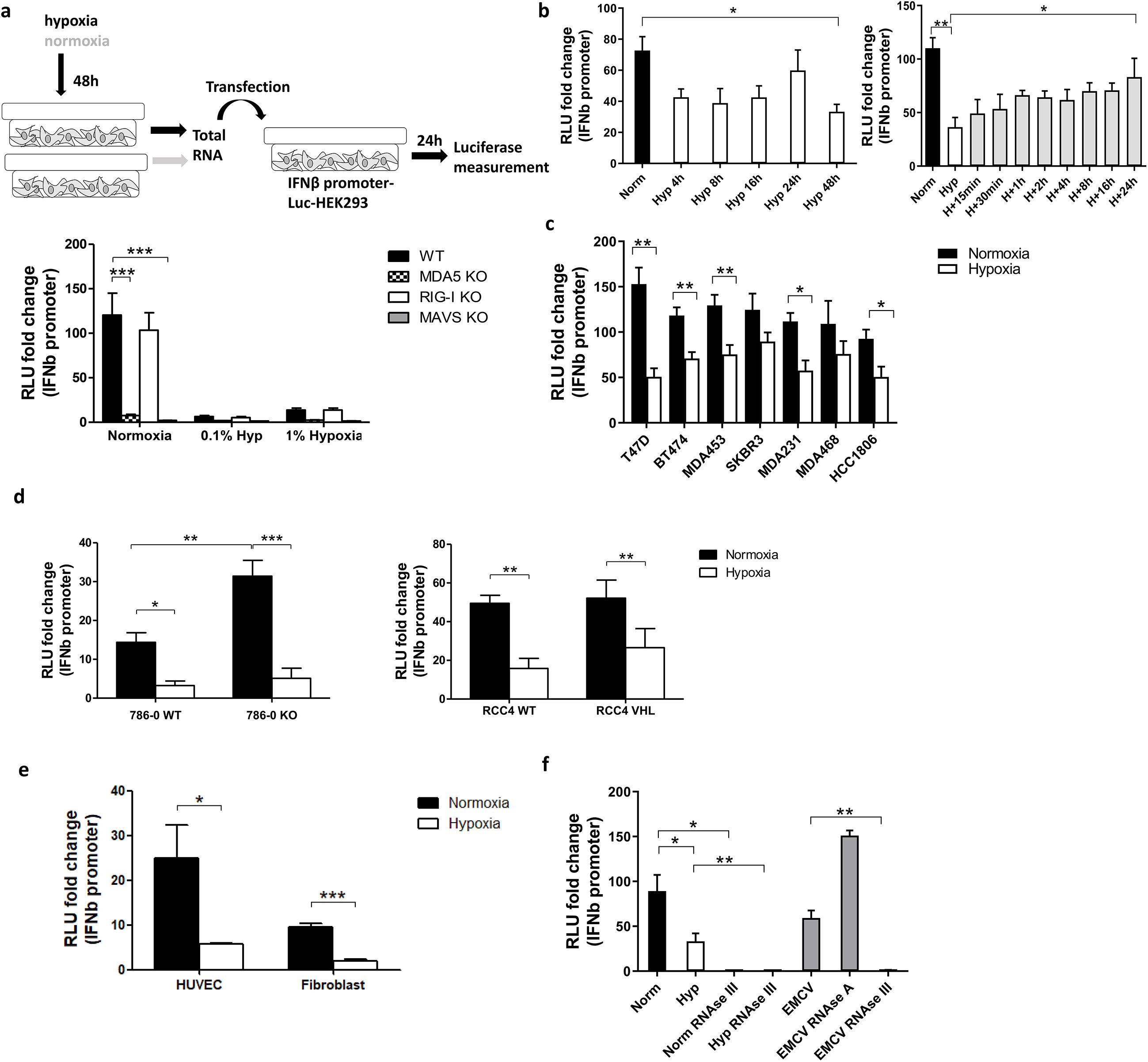
Hypoxia decreased *IFNβ* promoter stimulation. a) An outline of the experiment is shown (top, please see text for detail). IFN□ promoter reporter cells of the indicated genotypes were transfected with total RNA extracted from MCF7 cells exposed for 48h to normoxia, 1% hypoxia and 0.1% hypoxia. 24h after transfection, reporter cells were lysed and firefly luciferase activity was determined (bar charts; RLU, relative light units). Data from control cells treated with transfection reagent only were used to calculate RLU fold changes after RNA transfection (n=10). b) *IFNβ* promoter stimulation time course using RNA from MCF7 cells exposed to normoxia or 0.1% hypoxia for 4h, 8h, 16h, 24h and 48h (left panel, n=3), and reoxygenation for 15min, 30min, 1h, 2h, 4h, 8h, 16h and 24h after 48h in 0.1% hypoxia (right panel, n=3). c) *IFNβ* promoter stimulation using RNA from a panel of breast cancer cell lines exposed to normoxia or 0.1% hypoxia for 48h. d) *IFNβ* promoter stimulation using RNA from 786-0 WT or 786-0 HIF2α KO cells (786-0 KO, left panel, n=3), and RCC4 EV or RCC4 VHL (right panel, n=3) exposed to normoxia or 0.1% hypoxia for 48h. e) *IFNβ* promoter stimulation using RNA from non cancerous cell lines exposed to normoxia or 0.1% hypoxia for 48h (n=3). f) *IFNβ* promoter stimulation using RNA from MCF7 cells treated with RNAse A or RNAse III, and EMCV dsRNA as positive control (n=3). Number of replicates indicate biological replicates and data is shown as mean±SEM. * p<0.05, ** p<0.01, *** p<0.001.

As 0.1% hypoxia had a greater effect than 1% hypoxia (fig. 1a), subsequent experiments were performed under 0.1% hypoxic conditions. A time course in hypoxia for 4h, 8h, 16h, 24h and 48h showed that 4h in hypoxia was enough to lower IFNβ promoter stimulation and it was maintained up till 48h (fig. 1b left panel). The time course for recovery after reoxygenation following 48h hypoxia was evaluated at 15min, 30min, 1h, 2h, 4h, 8h, 16h and 24h. Reoxygenation caused a gradual recovery of IFNβ promoter stimulation reaching normoxic basal levels at 24h (fig. 1b right panel).

A panel of breast cancer cell lines with different receptor status were used to rule out a cell line dependent effect of hypoxia in MCF7 cells. Hypoxic RNA was much less effective than normoxic RNA in stimulating IFNβ promoter in all cell lines (fig. 1c).

### Hypoxic reduction of dsRNA formation is HIF1/2α independent

Normoxic and hypoxic RNA from 786-0 WT (HIF1α deficient and HIF2α upregulated due to *VHL* mutation) and 786-0 HIF2α KO (hereafter 786-0 KO, HIF1α and HIF2α deficient), and RCC4 WT (*VHL* mutation leading to HIF1α and HIF2α constitutive overexpression) and RCC4 VHL (*VHL* restored causing HIF1α and HIF2α downregulation) was tested. In both cell lines and all genotypes, hypoxic RNA triggered significantly lower *IFNβ* promoter stimulation (fig. 1d), highlighting the HIF-independent effect. However, 786-0 KO cells showed higher *IFNβ* promoter activation under normoxia compared to 786-0 WT, suggesting an effect of HIF2α in suppressing IFNβ induction, although minimal compared to the effect of hypoxia.

We also tested normal endothelial cells (HUVECS) and fibroblasts. Again, hypoxic RNA significantly reduced the activation of *IFNβ* promoter (fig. 1e) pointing to a general effect of hypoxia independently of cancer.

To analyse which RNA species from the total RNA were responsible for the *IFNβ* promoter induction, normoxic and hypoxic RNA from MCF7 cells was treated with different RNAses and EMCV (*Encephalomyocarditis Virus*), containing only dsRNA, was used as positive control. RNAse III (specific for dsRNA) treatment completely abolished normoxic, hypoxic and EMCV RNA induced *IFNβ* promoter activity, whereas RNAse A (specific for single-stranded RNA, ssRNA) treatment did not affect the luciferase signal (fig. 1f), showing that endogenous dsRNAs are responsible for the *IFNβ* promoter activation rather than ssRNA.

### Imaging of dsRNA levels downregulation under hypoxia

To visualise the downregulation of dsRNA levels in hypoxia, fluorescence microscopy was performed using the J2 antibody which is widely used to specifically detect dsRNA^18,21^. There was substantial variability of dsRNA intensity among individual cells, but dsRNA staining was significantly lower in MCF7 hypoxic cells (fig. 2a). The downregulation was time-dependent and significant after 16h in hypoxia (fig. 2b). To confirm the HIF1α/HIF2α-independence observed in the *IFNβ* promoter assay, 786-0 WT and 786-0 KO cells were stained and both cell lines showed significantly lower dsRNA levels in hypoxia (fig. 2c).

**Figure 2.**
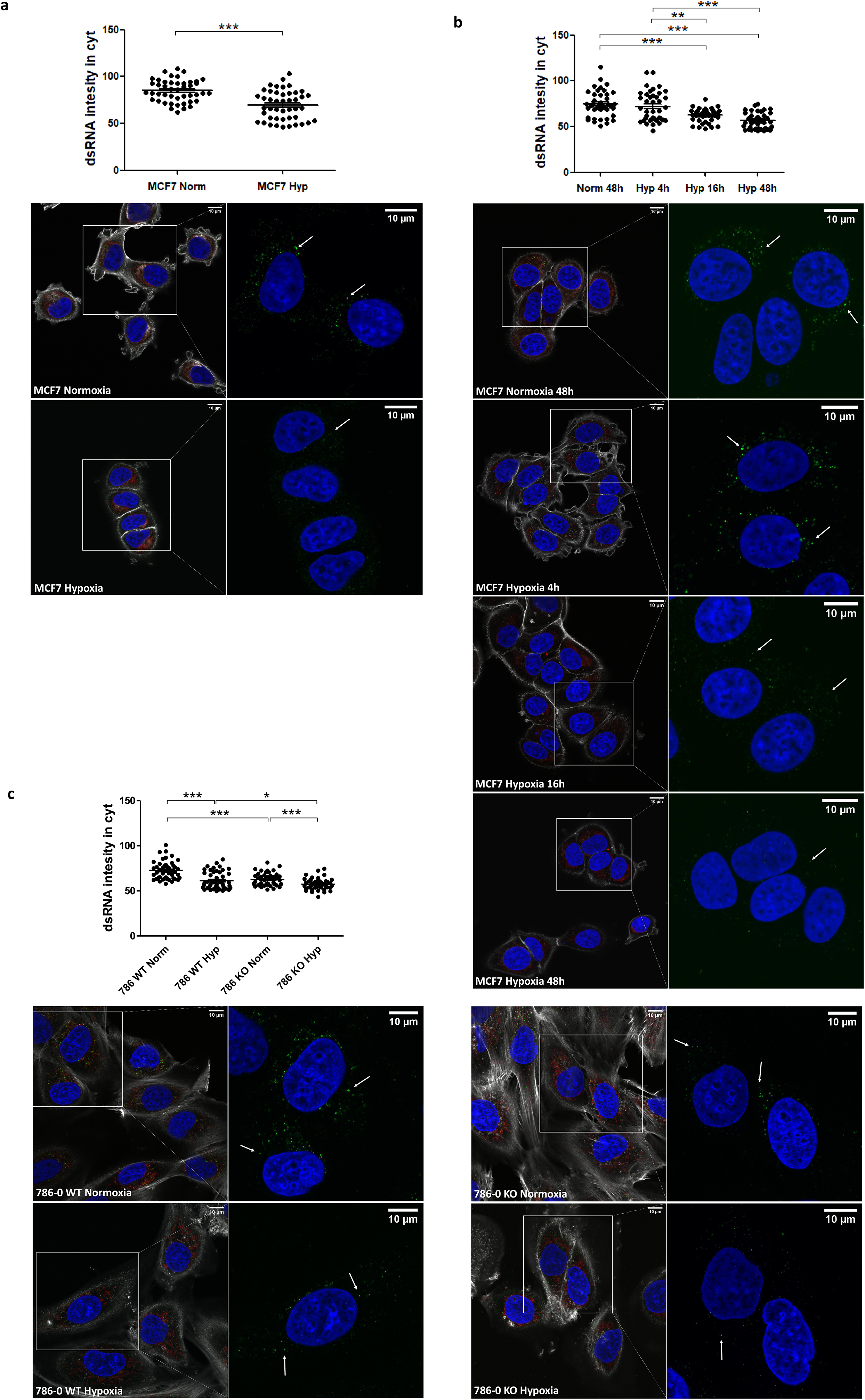
dsRNA staining is significantly lower under hypoxia independently of HIF1α/2α expression. a) dsRNA was stained using J2 antibody in MCF7 cells exposed to normoxia (n=45 cells) or 0.1% hypoxia (n=45 cells) for 48h from 3 independent replicates. b) dsRNA was monitored during a time course of MCF7 cells in normoxia (n=40 cells), or exposed to 0.1% for 4h (n=40 cells), 16h (n=40 cells), and 48h (n=40 cells) from 3 independent replicates. c) HIF1α/2α involvement was evaluated by staining dsRNA in 786-0 WT cells (786 WT, n=46 normoxic cells and n=46 hypoxic cells), and 786-0 HIF2α-KO cells (786 KO, n= 45 normoxic cells and n=45 hypoxic cells) from 3 independent replicates. Data is shown as mean±SEM. * p<0.05, ** p<0.01, *** p<0.001. Green: J2 antibody staining, blue: DAPI, and red: mitotracker staining. Scale bars correspond to 10μm.

### mtDNA and effect of mutation status on dsRNA in hypoxia

It was recently shown that 99% of endogenous dsRNA originates during mtDNA transcription^18^. Therefore, we tested cells lacking mtDNA (Rho Zero) and found significantly lower dsRNA staining in the Rho Zero cells than the parental 143B cell line (fig. 3a) and similar reduction in *IFNβ* promoter activation (fig. 3b). This strongly suggested that regulation of mitochondria was a key mechanism for hypoxic downregulation of the dsRNA. Surprisingly, there was no difference between normoxia and hypoxia in the 143B parental cell line, either in the dsRNA staining or in the *IFNβ* promoter assay. This was the only cell line among all tested in which hypoxia did not downregulate the type I IFN pathway.

**Figure 3.**
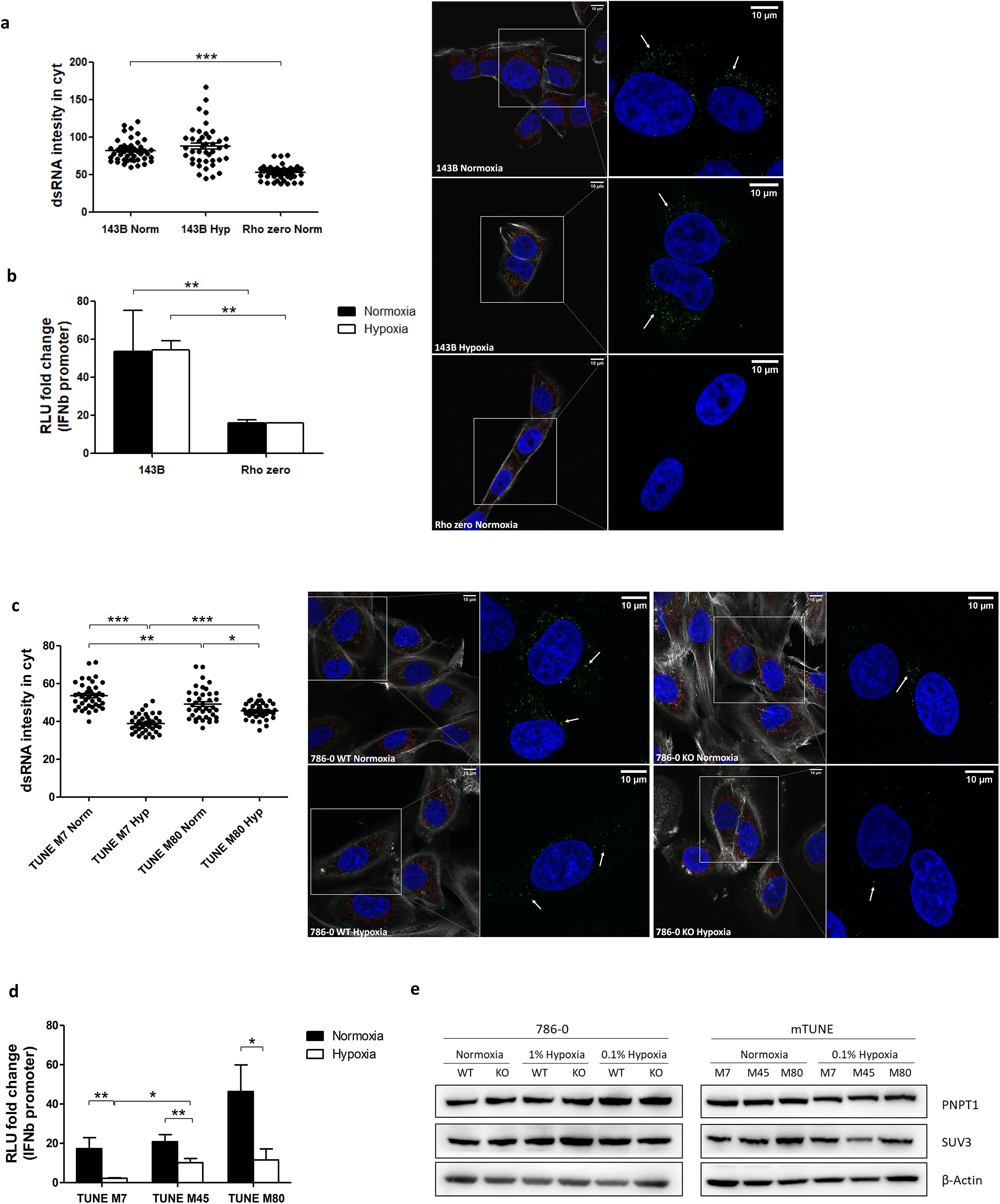
Mitochondrial alterations did not affect dsRNA staining reduction under hypoxia. a) Representative images showing dsRNA staining using J2 antibody in 143B WT cells in normoxia (n=44 cells) or 0.1% hypoxia (n=42 cells) for 48h and in 143B lacking mtDNA (Rho Zero) exposed to normoxia (n=40 cells) from 3 independent replicates. b) *IFNβ* promoter stimulation was evaluated as explained in fig.1a using RNA from 143B WT and Rho Zero cells cultured in normoxia and 0.1% hypoxia for 48h (n=3; RLU, relative light units). c) Representative images showing dsRNA staining in U2OS isogenic lines harbouring 7% vs 80% of heteroplasmy for the mtDNA mutation m8993T>G (mTUNE M7 normoxia n=40 cells and 0.1% hypoxia n=40 cells, M80 normoxia n=40 cells and 0.1% hypoxia n=40 cells) from 3 independent replicates. d) *IFNβ* promoter stimulation using RNA from mTUNE M7, M45 and M80 cells in normoxia and 0.1% hypoxia for 48h (n=3). e) Western blot showing PNPT1 and SUV3 dsRNA degrading enzymes protein levels in normoxia and hypoxia (n=3). Number of replicates indicate biological replicates and data is shown as mean±SEM. * p<0.05, ** p<0.01, *** p<0.001. Green: J2 antibody staining, blue: DAPI, and red: mitotracker staining. Scale bars correspond to 10μm.

Differences in metabolism in hypoxic mitochondria were considered a potential contribution to dsRNA release. In hypoxia there is a shift to reductive carboxylation for glutamine utilisation by mitochondria^22^ and to test this we used U2OS cells harbouring different degrees of heteroplasmy for the mtDNA mutation m8993T>G ranging from 7 to 80% (U2OS mTUNE M7, M45, M80)^23^. The basal level of cytoplasmic dsRNA was similar in M7 and M45 (data not shown), as were their metabolic profiles^23^, but higher in M80 cells. However, hypoxia downregulated dsRNA levels in all mTUNE cell lines independently of the mutation level (fig. 3c) and caused significantly lower *IFNβ* promoter activation (fig. 3d).

Moreover, it was previously reported that mtdsRNA accumulated and triggered the type I IFN pathway when the degrading enzymes were inhibited (PNPT1, SUV3)^18^. However, neither PNPT1 nor SUV3 protein levels were affected by 0.1% hypoxia for 48h (fig. 3e).

### Hypoxia reduces mtDNA transcription and mitochondrial ribosomal protein (MRP) expression

We measured the expression of some mitochondrial encoded genes (*12S*, *ND3*, *ATP6* and *CYTB*) and also nuclear encoded genes involved in mtDNA transcription (*SHMT2* [mitochondrial serine hydroxymethyltransferase involved in the first step of the mitochondrial one-carbon metabolism cleaving serine to glycine], *POLRMT* [mitochondrial RNA polymerase that catalyses mtDNA transcription], *TFAM* [which stabilizes mtDNA, regulates mtDNA transcription, and is required for efficient promoter recognition by POLRMT] and *TFB1M* [mitochondrial dimethyladenosine transferase 1 whose interaction with POLRMT and TFAM is required for mtDNA transcription]). Both sets of genes were significantly downregulated in MCF7 cells cultured under 0.1% hypoxia (fig. 4a). Most of the tested genes showed lower expression after only 4h under hypoxia (*SHMT2*, *POLRMT*, *TFAM*, *ND3*, *ATP6* and *CYTB*) but it was significant for all when cultured for 16h in hypoxia. These results were confirmed by the general downregulation observed in mitochondrial encoded genes and nuclear encoded genes involved in mitochondrial function (from MitoCarta 2.0^24^) using RNA-seq data from MCF7 cells cultured in normoxia or 0.1% hypoxia for 48h (fig. 4b). As expected, nuclear encoded genes involved in glycolytic metabolism such as pyruvate dehydrogenase kinase 1 (*PDK1*) or glyceraldehyde-3-phosphate dehydrogenase (*GAPDH*) were upregulated in hypoxia.

**Figure 4.**
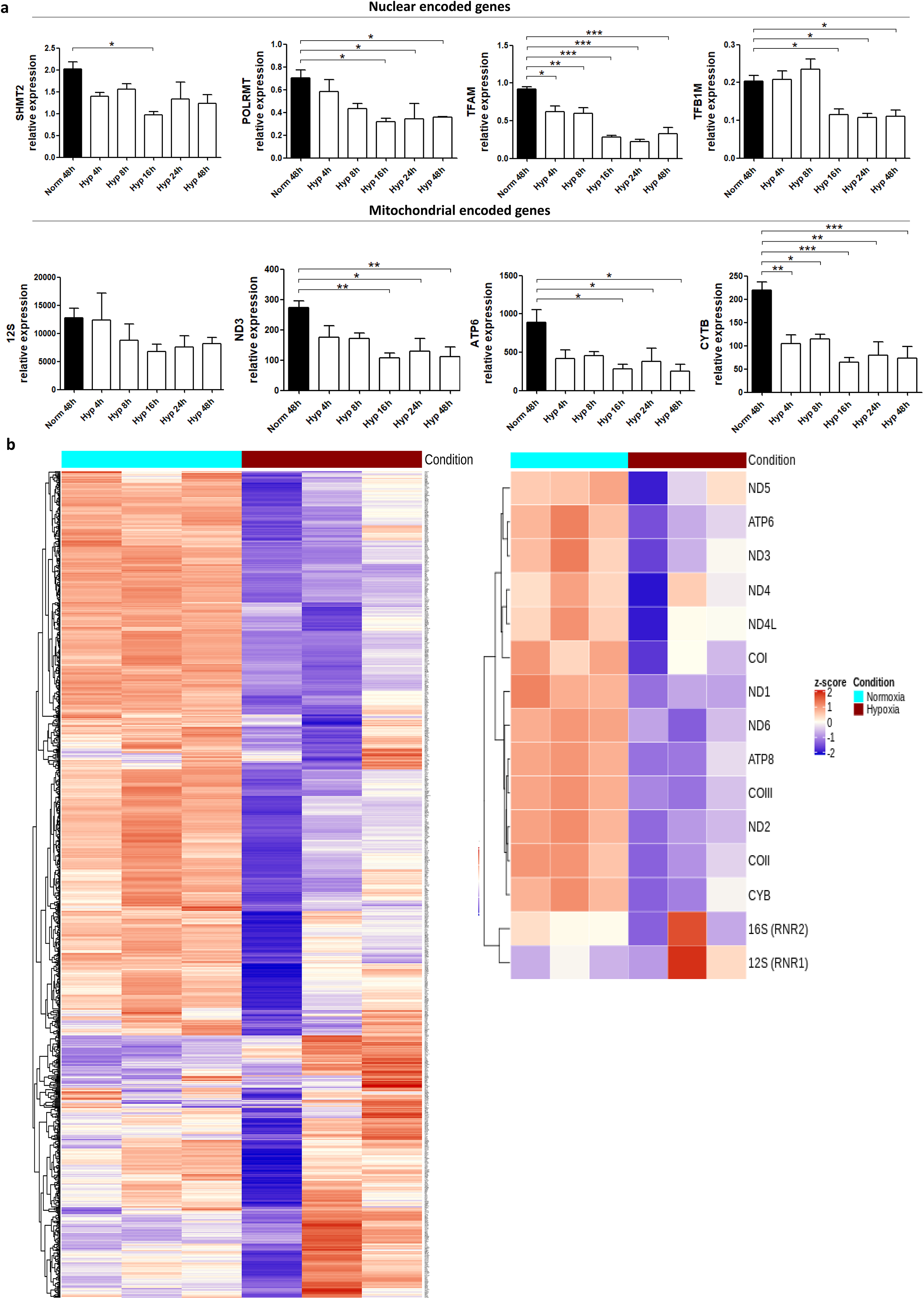
Hypoxia downregulates the expression of mitochondrial genes. a) RNA expression of mitochondrial encoded genes (*12S, ND3, ATP6, CYTB*) or nuclear encoded genes involved in mitochondrial function (*SHMT2, POLRMT, TFAM, TFB1M*) in MCF7 cells cultured in normoxia or 0.1% hypoxia for 4h, 8h, 16h, 24h and 48h was evaluated by qPCR (n=3). b) Heatmap showing expression of mitochondrial encoded genes (right panel) or 1158 nuclear encoded genes involved in mitochondrial function (left panel, from MitoCarta 2.0) in MCF7 cells cultured in normoxia or 0.1% hypoxia for 48h (n=3) from RNAseq experiment described in Materials section. Number of replicates indicate biological replicates and data is shown as mean±SEM.* p<0.05, ** p<0.01, *** p<0.001

We also determined the expression of *POLRMT, TFAM* and *TFB1M* and some mitochondrial encoded genes in the parental 143B and Rho zero cells. *12S, ND3* and *ATP6* mitochondrial genes were absent in Rho Zero cells whereas *POLRMT, TFAM* and *TFB1M* were expressed as previously reported^25^. However, hypoxia did not show any effect in the parental 143B cell line (Supplementary fig. 1a). This is potentially related to its cytosolic thymidine kinase (TK1) deficiency. Mitochondrial thymidine kinase (TK2) is not cell cycle regulated^26^. As the cytosolic and mitochondrial thymidine triphosphates are in rapid equilibrium and mainly produced by TK1^27^, it is possible that there is a more steady state of mtDNA replication, with a stable source of nucleotides from one compartment, which is not cell cycle dependent, in TK1 deficient cells. It was also reported that hypoxia decreased protein expression of mitochondrial ribosomal proteins (MRPs) involved in mtRNA translation^28^. We tested the expression of the mRNA coding for some of the MRPs under hypoxia in MCF7 cells and found that 16h of hypoxia significantly decreased their expression (fig. 5a). The same trend was observed in 786-0 WT and 786-0 KO cells although it was not always statistically significant (fig. 5b) pointing to a HIF1α/HIF2α-independent mechanism.

**Figure 5.**
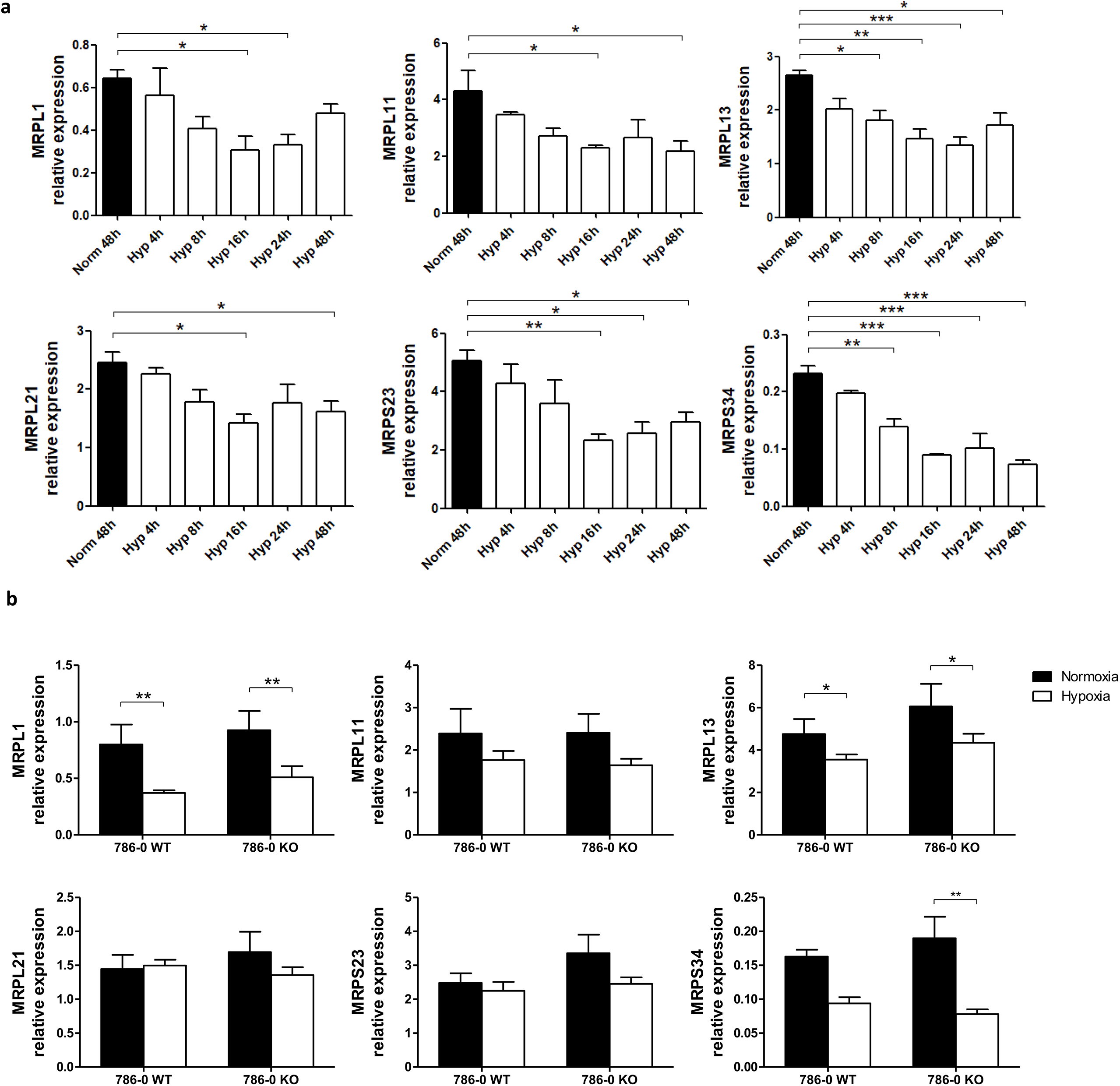
Expression of mitochondrial ribosomal proteins (MRPs) is downregulated under hypoxia. a) RNA expression of MRPs involved in mtRNA translation in MCF7 cells cultured in normoxia or 0.1% hypoxia for 4h, 8h, 16h, 24h and 48h was evaluated by qPCR (n=3). b) MRPs mRNA expression in 786-0 WT and 786-0 KO cells cultured in normoxia or 0.1% hypoxia for 48h was also evaluated by qPCR (n=3). Number of replicates indicate biological replicates and data is shown as mean±SEM. * p<0.05, ** p<0.01, *** p<0.001

Altogether, these data suggest that hypoxia leads to lower mtDNA transcription and thus lower production of dsRNA available to trigger the type I IFN response.

### mtRNA is the responsible for the *IFNβ* promoter induction

To assess the role of mtdsRNA in inducing the type I IFN pathway, MCF7 cells were cultured in normoxia or 0.1% hypoxia for 48h and their intact mitochondria were isolated. RNA from the mitochondrial and cytosolic fractions was extracted. Firstly, we confirmed that mitochondria were successfully isolated by analysing the expression of several nuclear and mitochondria encoded genes in both fractions (fig. 6a). Nuclear gene expression was detected both in the cytosolic and mitochondrial fractions, suggesting that the mitochondrial fraction could be slightly contaminated with mRNA from the cytoplasm. Nevertheless, the expression of mitochondrial encoded genes was significantly higher in the mitochondrial fraction. Interestingly, mitochondrial encoded genes were downregulated by hypoxia in the mitochondrial fraction but not affected in the cytosolic fraction (fig. 6a).

**Figure 6.**
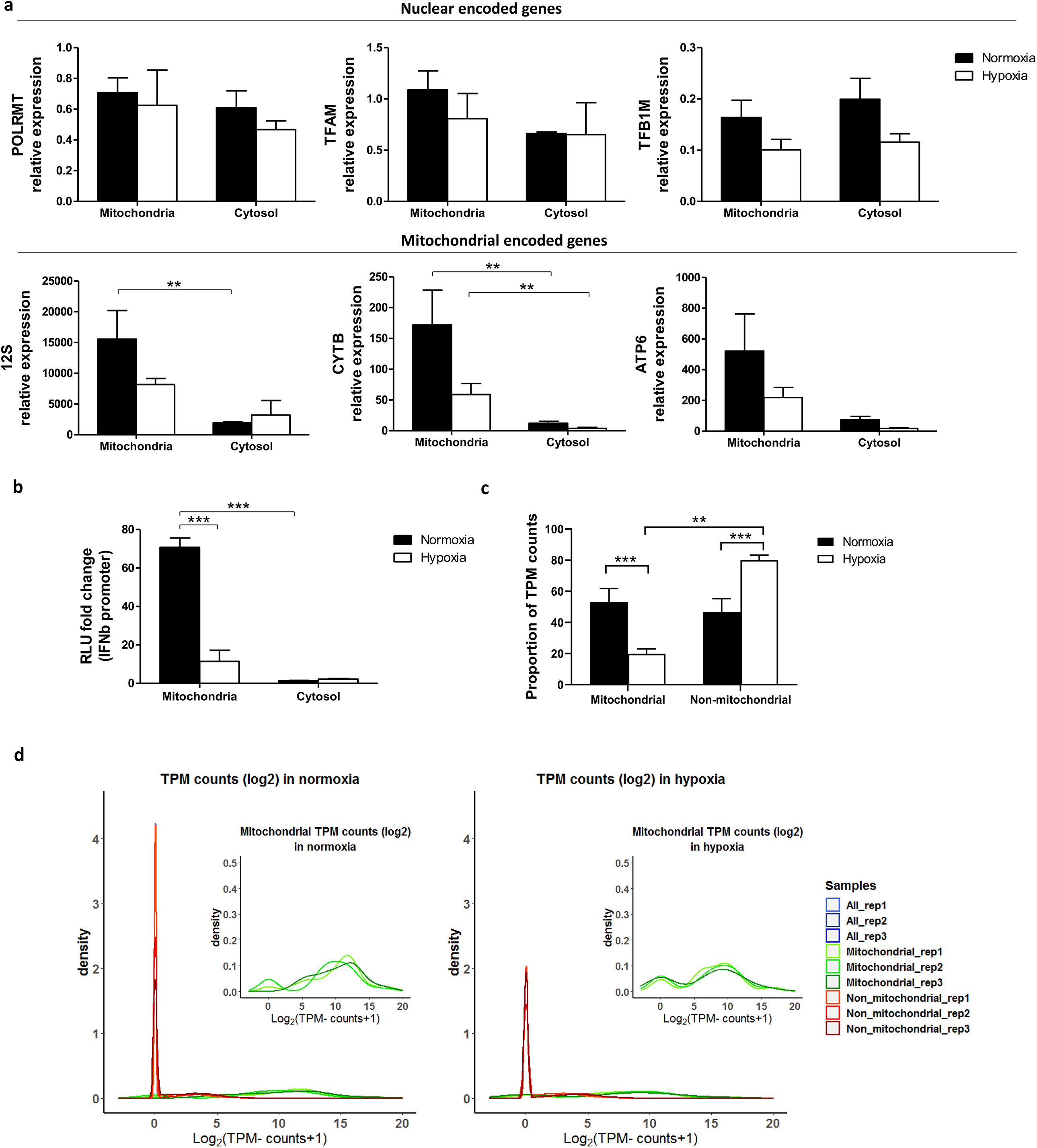
Mitochondrial RNA is responsible for *IFNβ* promoter activation. a) MCF7 cells cultured in normoxia and 0.1% hypoxia for 48h were fractionated. RNA expression of mitochondrial (*12S, CYTB, ATP6*) and nuclear encoded genes (*POLRMT, TFAM, TFB1M*) in mitochondrial and cytosolic fractions is shown by qPCR (n=3). Note difference in axes for mitochondrial versus nuclear genes. b) RNA from a) was used to activate the *IFNβ* promoter as described in fig. 1a (n=3; RLU, relative light units). c) dsRNA pull down experiment using J2 antibody in MCF7 cells exposed to normoxia or 0.1% hypoxia for 48h was performed. Bar graph shows proportion of transcripts per million (TPM) mitochondrial and non-mitochondrial reads (n=3). d) Density plots show mitochondrial (green), non-mitochondrial (red) and all reads (blue) in each replicate (rep) obtained from the dsRNA pull-down experiment in b) (n=3). Number of replicates indicate biological replicates and data is shown as mean±SEM. * p<0.05, ** p<0.01, *** p<0.001

The *IFNβ* promoter assay was then performed using these mtRNA and cytosolic RNAs. mtRNA induced *IFNβ* promoter stimulation, whereas the luciferase signal using cytosolic RNA was hardly detected. Importantly, hypoxic mtRNA caused significantly lower stimulation of *IFNβ* promoter (fig. 6b).

### dsRNA-enriched fraction in hypoxia showed lower mtRNA content

To assess the composition of the dsRNA pool in hypoxia, dsRNA pull-down was performed using the J2 antibody in MCF7 cells exposed to normoxia or 0.1% hypoxia for 48h, and the resultant RNA was sequenced. Reads were normalised as transcript per million (TPM). Interestingly, the percentage of mitochondrial reads was significantly lower in hypoxia than in normoxia (fig. 6c), and 22 out of 37 mitochondrial encoded genes were significantly downregulated in hypoxia (Supplementary table 1). Density plots in normoxia and hypoxia showed that the non-mitochondrial genes pulled-down by J2 antibody had few reads which probably correspond to background noise (red peak, fig. 6d).

### Lower mtRNA in hypoxia is not due to increase mitophagy

To evaluate whether the downregulation of mtdsRNA in hypoxia was due mitophagy, we knocked down one of the main genes involved in this process, *BNIP3* (BCL2 Interacting Protein 3) in MCF7 exposed to 0.1% hypoxia for 48h (siBNIP3 and control siRNA, siCON). In the knock-down cells, BNIP3 showed no induction in hypoxia, either at mRNA or protein level. The expression of various ISGs (*DDX58*, *MX1*, *IFIT1*, *IFIT2*, *ADAR-p150* and *ISG15*) was tested. Hypoxia downregulated ISG expression^19^, but no differences were found between siCON and siBNIP3 in hypoxia (fig. 7b), suggesting that hypoxic upregulation of BNIP3 does not interfere in the type I IFN signalling. This was further confirmed by the *IFNβ* promoter assay, in which hypoxia-induced inhibition on the *IFNβ* promoter activation was not recovered upon BNIP3 silencing (fig. 7c).

**Figure 7.**
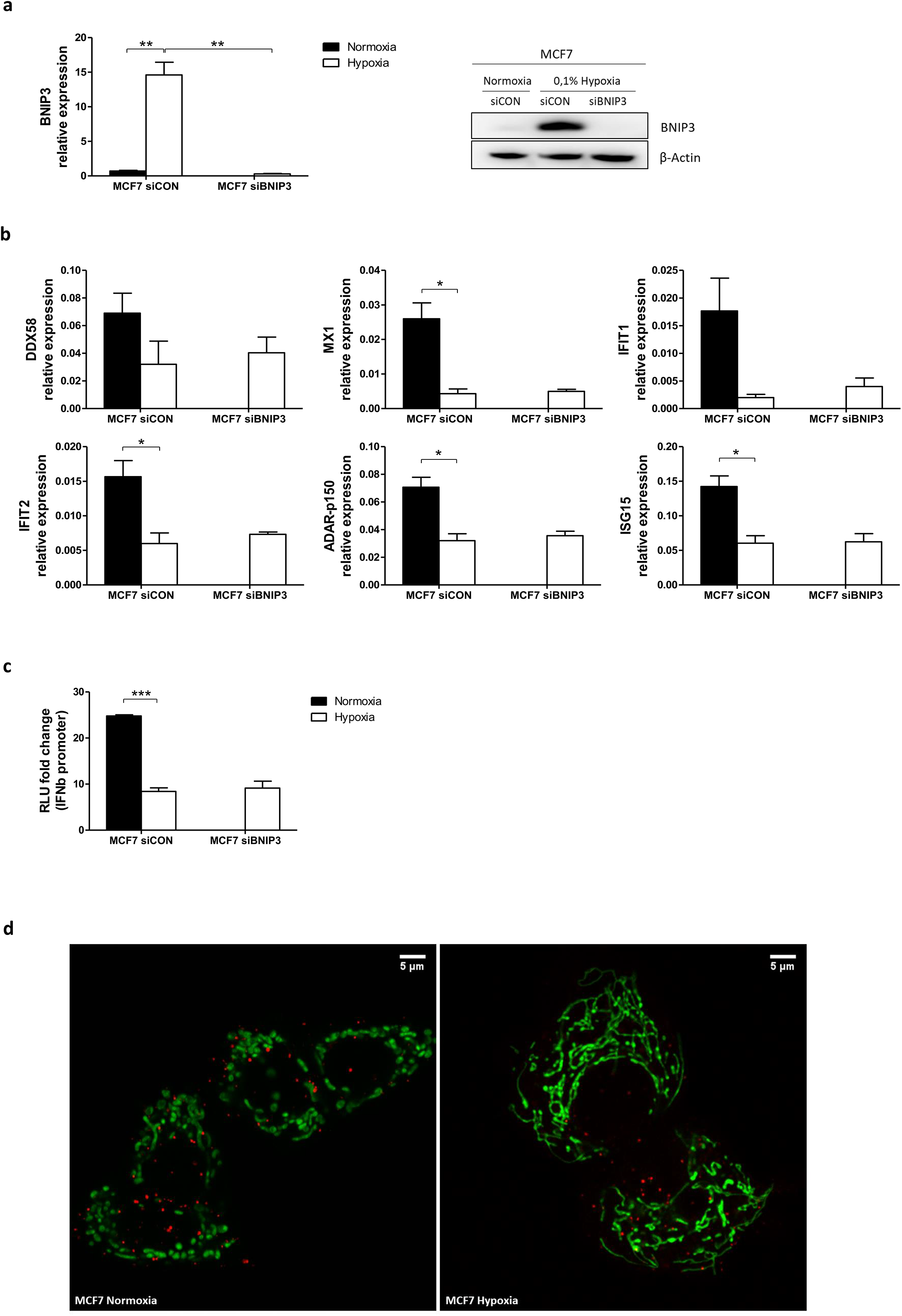
Mitophagy is not involved in mtdsRNA reduction under hypoxia. a) Silencing of BNIP3 was confirmed by qPCR (left panel) and western blot (right panel) in siBNIP3 MCF7 cells in hypoxia vs control (siCON) (n=3). b) RNA expression of IFN-induced genes (ISGs) in siBNIP3 MCF7 cells vs siCON was performed by qPCR (n=3). c) RNA from a) was used to evaluate *IFNβ* promoter activation after BNIP3 silencing (n=3; RLU, relative light units). d) Representative image showing lack of colocalization between the mitochondrial (green) and lysosomal (red) markers in MCF7 cells exposed to normoxia or 0.1% hypoxia for 48h (n=3). Number of replicates indicate biological replicates and data is shown as mean±SEM. * p<0.05, ** p<0.01, *** p<0.001

Live cell immunofluorescence was performed using mitochondrial and lysosomal markers in MCF7 cells cultured in normoxia or 0.1% hypoxia for 48h. As previously reported^29^, mitochondria clearly exhibited morphological changes under low oxygen conditions and appeared more elongated and located closer to the nucleus (fig 7d). However, no colocalization of the mitochondrial and lysosomal markers was observed in normoxia or hypoxia, suggesting that mitophagy is not increased in hypoxia and does not decrease mtdsRNA.

### Effect of mitochondria-targeting drugs in dsRNA levels

As mitochondria are the main source of dsRNA in the cells^18^, the ability of different drugs targeting mitochondria to affect dsRNA levels, as a therapy to overcome hypoxic effects, was assessed in MCF7 cells. Briefly, the drugs used were: chloramphenicol which inhibits mitochondrial protein synthesis via binding to the 50S subunit of the 70S mitochondrial ribosomes; ABT-737 induces cell apoptosis by inhibiting the anti-apoptotic molecule BCL2, as well as mitophagy; G-TPP accumulates in the mitochondria of tumour cells and by inhibiting the heat shock protein 90 (Hsp90) promotes cell apoptosis; mubritinib and metformin decrease mitochondrial respiration by inhibiting ETC complex I; and the Vps34 inhibitor SAR405 alters vesicle trafficking and inhibits autophagy by blocking autophagosome formation.

Mubritinib and G-TPP treatment decreased *IFNβ* promoter activation under normoxia by 4- and 2-fold, respectively (supplementary fig. 2). Furthermore, hypoxia was not able to further inhibit *IFNβ* promoter activity in MCF7 cells treated with these two drugs.

Thus, these mitochondrial targeting drugs did not result in increased dsRNA, and although mubritinib and G-TPP inhibited *IFNβ* promoter stimulation, hypoxia showed a greater effect.

### Tissue distribution of immunostimulatory RNA

Total RNA samples from different human tissues were obtained to assess their effects in the *IFNβ* promoter reporter assay. RNA from some tissues including brain, heart, kidney and testis strongly induced the *IFNβ* promoter reporter (fig. 8a). In contrast, RNA from other tissues including skeletal muscle and pancreas had little effect. ISG expression was also determined in RNA samples from some tissues. *MX1, IFIT1* and *IFNB1* exhibited much higher expression in testis and brain than in skeletal muscle (fig. 8b), confirming the results from the *IFNβ* promoter assay.

**Figure 8.**
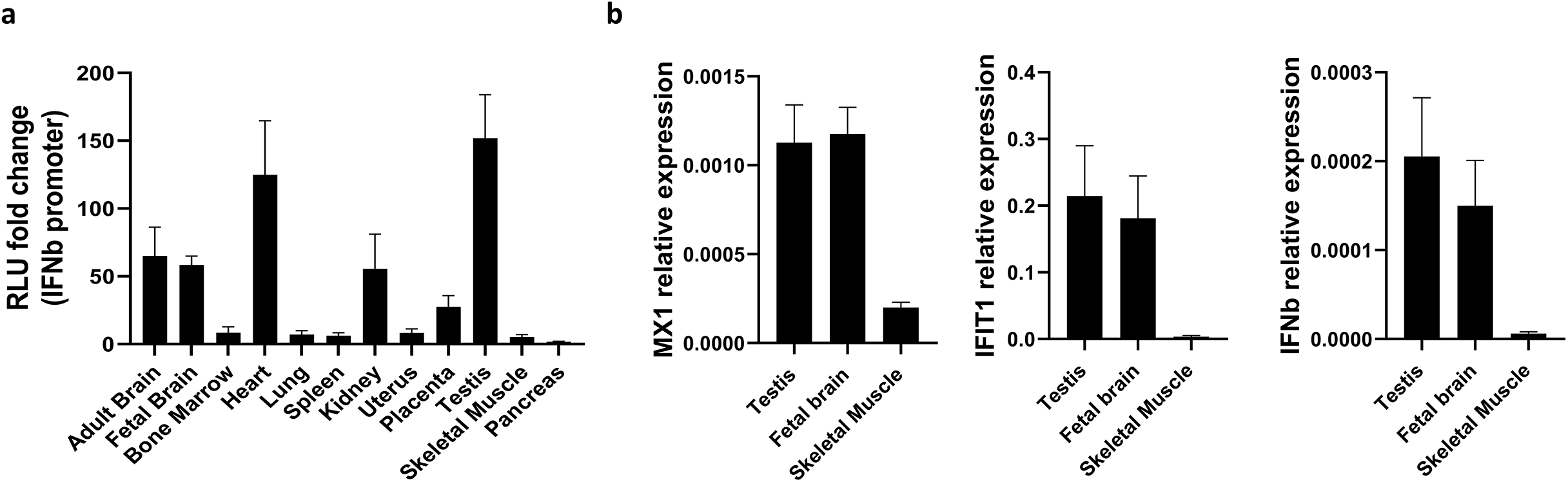
Level of type I IFN pathway activation is tissue-dependent. a) The *IFNβ* promoter reporter assay shown in Fig. 1a was used to assess the immunostimulatory properties of RNA samples from different tissues (n=3). b) qPCR showing expression of *MX1, IFIT1* and *IFNβ* in different tissues from a) (n=3). Number of replicates indicate biological replicates and data is shown as mean±SEM.

## DISCUSSION

In this paper we have shown that hypoxia caused less formation of mtdsRNA in different cancer and normal cell lines leading to lower activation of *IFNβ* promoter. This repressive effect was HIF1α/2α-independent as hypoxia led to downregulation of mitochondrial gene expression in 786-0 HIF2α-KO cells, as in most other cell lines. However, 786-0 WT cells did not show lower mitochondrial gene expression in hypoxia. Those cells express only HIF2α, which enhances c-Myc transcriptional activity^30^ and c-Myc increases the expression of POLRMT, thus increasing mtDNA transcription^31^. Possibly this could compensate for the effect of hypoxia, and other transcription factors such as c-Jun and NF-κB are noted to have a mitochondrial localisation and can be regulated in hypoxia in a HIF-independent manner^32^.

Endogenous mtRNA and, specifically mtdsRNA, was responsible for triggering the *IFNβ* promoter activation via the MDA5/MAVS and not RIG-I/MAVS sensing pathway and the mechanism of reduction in hypoxia was investigated. It was recently reported that 99% of endogenous dsRNA was produced as a consequence of mtDNA transcription, and inhibition of the mtdsRNA degrading enzymes SUV3 and PNPT1 increased type I IFN signalling^18^. However, hypoxia did not affect the expression of these degrading enzymes, thus suggesting that lower mtdsRNA presence in hypoxia could be a consequence of lower mitochondrial transcription under low oxygen conditions rather than higher degradation.

dsRNA pull-down experiments showed that most of the reads corresponded to the mitochondrial chromosome. However, the percentage of mitochondrial reads was significantly lower in hypoxia and the expression of mitochondrial encoded genes was significantly downregulated. Supporting this hypothesis, we found decreased expression of mitochondrial encoded genes and nuclear encoded genes involved in mitochondrial transcription and mitochondrial ribosomal proteins when cells were cultured for only 4h under hypoxia and it became significant at 16h. Moreover, dsRNA staining was also significantly lower at 16h under hypoxia.

Hypoxic-induced mitophagy via BNIP3 was another possible explanation for decreased dsRNA formation^33,34^. However, our data showed that BNIP3 silencing did not revert the hypoxia-caused downregulation of several ISGs or *IFNβ* promoter activation. Moreover, live cell imaging of mitochondria and lysosomes supported this result, as no colocalization of mitochondrial and lysosomal markers was observed either in normoxia or hypoxia.

Moreover, we investigated mitochondria-targeting drugs as potential regulators of dsRNA release. Low oxygen concentrations lowered the *IFNβ* promoter activation to the same extent as controls for most of the mitochondria-targeting drugs and those inhibitors did not affect normoxic levels. Inhibition of respiration by metformin, a complex I inhibitor, did not affect *IFNβ* promoter activation. In contrast, mubritinib and G-TPP, caused a downregulation that reached hypoxic levels, with no further effect of hypoxia. However, mubritinib, which also targets complex I^35^, is known to inhibit HER2^36^. HER2 inhibition by trastuzumab increased the expression of pro-apoptotic proteins, which induced the opening of mitochondrial permeability transition pores, ROS production and mitochondrial dysfunction due to loss of mitochondrial membrane potential^37,38^. Therefore, mubritinib could further contribute to mitochondria dysfunction and the consequent decrease in dsRNA synthesis in MCF7 cells, which have low level of functional HER2^39,40^. On the other hand, G-TPP specifically inhibits the mitochondrial protein-folding chaperone Hsp90, generating unfolding protein stress in the mitochondria^41^ which could decrease gene transcription. G-TPP has also been described to induce mitophagy^42^, but as shown above, it is not likely to be the mechanism to decrease dsRNA. The similarity of reduction by these 2 inhibitors to that induced by hypoxia, and lack of further suppression suggest these pathways could overlap e.g. by inhibiting RNA synthesis.

Basal levels of dsRNA in different tissues without hypoxic stress was assessed using human total RNA, assuming that the assay measured only dsRNA. RNA from testis, brain, heart and kidney were strong activators of *IFNβ* promoter, and these tissues also showed higher expression of type I IFN genes (*MX1, IFIT1* and *IFNB1*). Mitochondrial mass is well correlated with citrate synthase and cytochrome oxidase activity, mtDNA copy number and mitochondrial gene expression, and all these parameters are greater in tissues with higher bioenergetics and metabolic demands such as heart^43^. Although skeletal muscle RNA hardly triggered *IFNβ* promoter activation whereas heart RNA was far more effective, this could be explained by low basal activity of mitochondria in resting striated muscle. Additionally, higher oxygen concentrations in tissues such as brain, heart and kidney could stimulate higher turnover^44^. Interestingly, some tumour types such as bladder, breast, esophageal, head and neck, kidney and liver showed significantly lower mtDNA content than paired adjacent normal tissue, and this was associated with lower patient survival^45^. Expression could be even lower in hypoxic areas having impact in anticancer therapies that rely on functional type I IFN signalling.

To sum up, we have shown that hypoxia caused significantly lower mtdsRNA production, probably due to a decrease in mitochondrial transcription rather than increased degradation, thus leading to lower activation of *IFNβ* promoter, and consequently to lower type I IFN response that could contribute to the immunosuppression observed in hypoxic environments and to chemotherapy and radiotherapy resistance.

## MATERIALS

### Human tissues, cell culture and transfection

Human total RNA used in this manuscript was purchased to Takara Bio/Clontech. MCF7, T47D, BT474, MDA-MB-231, MDA-MB-453, MDA-MB-468, RCC4, 786-0, U2OS mTUNE, and human fibroblasts were cultured in DMEM low glucose medium (1g/l; Thermo Fisher Scientific) supplemented with 10% FBS no longer than 20 passages. HUVEC cells were purchased to Lonza and grown in EGM2 medium for maximum 7 passages. They were all mycoplasma tested every 3 months and authenticated during the course of this project. Cells were subjected to 1% or 0.1% hypoxia for the periods specified in each experiment using an InVivO_2_ chamber (Baker).

U2OS mTUNE glioblastoma cell lines were kindly donated by Dr Christian Frezza. Three isogenic cell lines with different levels of heteroplasmy of mutated mtDNA were used: M7, M45 and M80^23^.

143B and 143B Rho Zero (Rho Zero) cells were a gift of Dr Karl Morten. Rho Zero cells were generated by treating 143B cells (TK1 deficient) with 10μM 2’,3’-dideoxycytidine (ddC) for 10 days. Both cell lines were grown in high glucose DMEM (Gibco) supplemented with 10% FBS. Rho Zero cell culture media was additionally supplemented with 50 μg/mL uridine (A15227.06, Alfa Aesar).

### Drug treatment

MCF7 cells were cultured in normoxia or 0.1% hypoxia and treated for 48h with the following mitochondria-targeting drugs: 5μM Vps34 inhibitor SAR405 (16979, Cayman Chemical), 200μM chloramphenicol (C0378, Sigma-Aldrich), 100nM mubritinib (S-2216, Selleckchem), 5μM ABT-737 (sc-207242, Santa Cruz Biotechonology), 2mM metformin (D150959, Sigma-Aldrich) or 5μM gamitrinib-triphenylphosphonium (G-TPP, HY-102007, MedChemExpress).

### siRNA transfection

BNIP3 siRNA transfection (supplementary table 2) was performed in Optimem reduced serum medium at a final concentration of 5nM, the following day cells were exposed to 0.1% hypoxia for 48h. siRNA control was done in parallel and the following day was subjected to normoxia or 0.1% hypoxia for 48h. Oligofectamine (12252-011, Thermo Fisher Scientific) was used following the manufacturer’s instructions.

### Western blot

Whole cell lysates were prepared with RIPA buffer (R0278, Sigma) containing protease (cOmplete, 11697498001) and phosphatase (phosSTOP, 4906845001) inhibitors. Samples were subjected to SDS-PAGE and transferred onto PVDF membranes (IPVH00010, Millipore), after blocking, membranes were incubated overnight with primary antibodies (supplementary table 3) at 4°C. They were later washed and incubated with HRP-anti-mouse/rabbit secondary antibodies (Gibco). Development was performed with Amersham ECL Prime Western Blotting Detection Reagent (GERPN2232, GE Healthcare Life Sciences) using ImageQuant™ LAS 4000. Stripping with Restore PLUS Western Blot Stripping Buffer (46430, Invitrogen) was performed to blot different antibodies in the same membrane.

### RT-qPCR

RNA was extracted using the Tri-Reagent protocol (T9424, Sigma) and 1μg was reverse transcribed with the High Capacity cDNA reverse transcription kit (44368813, Thermo Fisher Scientific) using random hexamer primers. The PCR reaction containing SensiMix™ SYBR Green^®^ No-ROX Kit (QT650-20, Bioline) was run on a 7900 Real time PCR System (Applied Biosystems) with standard cycling conditions: 10 minutes 95°C, and 40 cycles of 15 seconds 95°C followed by 1 minute 60°C. Gene expression was analysed with the Ct method using *HPRT1* expression for normalization. The primers used are listed in Supplementary table 4.

### *IFNβ* promoter reporter assay

HEK293T-P125 reporter cells (stably expressing the *IFN-β* promoter-Luciferase region^20^) were used to detect specifically immunostimulatory RNAs as they do not express cGAS and the expression of STING is very low. 4×10^4^ cells per well were seeded in 96-well plates. Next day, cells were pre-treated with 30U/ml of IFN-A/D (I4401, Sigma), and after 24h of incubation fresh medium was added and cells were transfected with 100ng of total RNAs from cell cultures or human tissues using Lipofectamine 2000^®^ (11668-019, Thermo Fisher Scientific). As positive controls, 1ng of IVT-RNA or V-EMCV-RNA was used^20^. 24h post-transfection, cell were lysed and measured using OneGlo luciferase assay (E6120, Promega) in a FluorOPTIMA luminometer.

### Immunofluorescence for dsRNA

Cells were plated on coverslips (VWR Collection) and exposed to 0.1% O_2_ hypoxia for 48h. Prior to fixation, mitochondria were stained for 1h with 200nM MitoTracker Deep Red (M22426, Thermo Fisher Scientific). After, cells were washed and fixed with 4% (v/v) paraformaldehyde (PFA) for 8 min at RT. Then, cells were washed and permeabilized with 0.1% Triton X-100 for 20 min at RT. PFA was neutralized with 0.1M glycine for 10 min at RT. After washing three times, cells were incubated for 60 min with blocking solution (PBS containing 1% (w/v) BSA and 10% (v/v) normal goat serum (ab7481, Abcam)). Cells were incubated overnight at 4°C in a humidified chamber with J2 primary antibody (10010200, Scicons) at 1:200, and rhodamine phalloidin (R415, Invitrogen) at 1:40 in block solution. Cells were washed three times and incubated with goat anti-mouse IgG Alexa Fluor 488 (R37120, Invitrogen) secondary antibody at 1:500 and Hoechst 33342 (H3570, Invitrogen) at 1:1000 concentration in block solution for 1h at RT. After washing three times with PBS, coverslips were mounted with Vectashield® Mounting Medium (H-1000, Vector Labs) and sealed with nail polish.

### Immunofluorescence dsRNA image analysis

Slides were imaged in a Zeiss 880 Inverted confocal microscope (Zeiss) using a 63x Plan-Apochromat objective. Laser properties, acquisition mode and detectors were manually adjusted for each experiment. Fiji Image J software was used for image analysis, using specific macros created by Dr. Ulrike Schulze and Dr. Dominic Waithe. A minimum of 40 cells per condition were analysed.

### Mitochondria extraction

Mitochondria were isolated from MCF7 cells seeded in normoxia or 0.1% hypoxia for 48h using Mitochondria Isolation Kit for Cultured Cells (89874, Thermo Fisher Scientific), and following manufacturer’s instructions. Mitochondrial RNA was extracted using Tri Reagent, following the protocol previously explained. 200uL of the cytosolic fraction were also used to extract RNA.

### Immunoprecipitation of dsRNA

Protein G Dynabeads (10004D, Invitrogen) were washed and resuspended in NET-2 buffer. 5μg of J2 antibody or mouse IgG2 (400201, BioLegends) were bound to 100μL of beads for 1h at RT on a thermoshaker. Conjugated beads were washed three times with NET-2 Buffer. 80–90% confluent MCF7 cells from 10cm^2^ plate (×2) were washed with 10 ml of cold PBS. Cells were scraped and transferred to a falcon and spun at 500*g* at 4°C, 5 min. Cell pellet from one 10cm^2^ plate was lysed in 1 ml of NP-40 lysis buffer and transferred to a tube and incubated on ice for 5 min. Following centrifugation at 17,000*g* at 4°C for 5 min, supernatant was carefully transferred to a new tube. Total RNA was harvested from 10% input lysate using Tri Reagent. For immunoprecipitation, lysate was supplemented with 10 units of RNase free TurboDNase (AM2238, Ambion) at 10mM MgCl_2_ per 1 mL of mix. 100μL of J2-Dynabeads was added to 1mL of above lysate and left for 1–2h at 4°C. Following magnetic separation, beads were washed twice with 1mL of high salt washing buffer (HSWB). Beads were transferred to a new tube with NET-2 buffer and washed twice with the same buffer. J2-bound dsRNA was extracted with Tri Reagent. The RNA samples were sent for sequencing. NET-2 buffer (50mM Tris-Cl, pH 7.4, 150mM NaCl, 1mM MgCl_2_, 0.5% NP-40), NP-40 lysis buffer (50mM Tris-Cl pH 7.4, 150mM NaCl, 5mM EDTA, 0.5% NP-40), high salt wash buffer (50mM Tris-Cl pH 7.4, 1M NaCl, 1mM EDTA, 1% NP-40, 0.5% DOC, 0.1% SDS).

### RNA-sequencing and data analysis

Libraries for paired end sequencing were prepared using standard Illumina protocol and sequencing was performed using Illumina NovoSeq 6000 sequencer at Wellcome Centre for Human Genetics. Raw reads were processed using FASTQC and Cutadapt and aligned to the genome using STAR. Normalised counts of nuclear encoded genes involved in mitochondrial function and mitochondrial encoded genes were used for generation of heatmaps. In order to estimate the proportion of counts that map to mitochondrial and non-mitochondrial genes; transcript per million (TPM) normalized value of transcripts encoded by mitochondrial and non mitochondrial genome in different replicates was calculated. Proportion of TPM counts in mitochondrial or non-mitochondrial fractions was calculated by dividing the sum of TPM counts in the fraction by total TPM counts.

### Statistical analysis

GraphPad Prism 8.0 statistical analysis software (GraphPad Software) was used. If not otherwise specified, all the experiments were performed in 3 biological triplicates. ANOVA or ANOVA on ranks was normally used to study one variable in more than 2 groups depending on if they follow a normal distribution or not respectively. When two means were compared, t-test was performed if samples followed a normal distribution or Mann-Whitney if there was not a normal distribution. When analysing the influence of two different independent variables on one dependent variable, 2-way ANOVA was applied. In the graphs, the error bars depict the standard error of the mean (SEM).

## DATA AVAILABILITY

RNA-seq data is available in the Gene Expression Omnibus (GSE153557).

## ACKNOWLEDGEMENTS

We thank Dr Karl Morten (University of Oxford, UK) for the 143B and Rho Zero cells, Dr Charles Lawrie (Biodonostia Instituto de Investigación Sanitaria, Spain) for the 786-0 WT and HIF2α-KO cells, and Dr Christian Frezza (University of Cambridge, UK) for the mTUNE cells.

## AUTHOR CONTRIBUTION

EA, AM, JR and ALH designed the experiments, analysed the data and wrote the manuscript. AM and EA performed the experiments. AGDJ developed the *IFNβ* promoter assay and performed this assay for fig. 1a and fig. 8a. US and DW developed the ImageJ macro to quantify J2 immunofluorescence. NP performed all RNA-seq and J2-IP analysis. This research was supported by funding from Cancer Research UK (ALH), Breast Cancer Research Foundation (ALH), and Breast Cancer Now (ALH, AM).

