## Supplementary information for "Hypoxia regulates endogenous double-stranded RNA production via reduced mitochondrial DNA transcription"

Supplementary table 2. siRNA sequences.

| siRNA | Sequence (5'-3') |
|---|---|
| siCON | ACGACACGCAGGUCGUCAUTT* |
| siBNIP3 | UCGCAGACACCACAAGAUA |
| | GAACUGCACUUCAGCAAUA |
| | GGAAAGAAGUUGAAAGCAU |
| | ACACGAGCGUCAUGAAGAA |

Supplementary table 3. Antibodies.

| Protein | Antibody | Company | Dilution |
|---|---|---|---|
| β-actin | A3854 | Sigma | 1:50000 WB |
| HIF1a | 610958 | BD Transduction Lab | 1:1000 WB |
| HIF2a | NB100-122 | Novus Biologicals | 1:500 WB |
| SUV3 | A303-055A | Bethyl Laboratories | 1:1000 WB |
| PNPT1 | Ab96176 | Abcam | 1:1000 WB |
| J2 | 10010500 | Scions | 1:200 IF |

Supplementary table 4. Primers.

| Primers | Sequence (5'-3') |
|---|---|
| HPRT1_F | TGACACTGGCAAAACAATGCA |
| HPRT1_R | GGTCCTTTTCACCAGCAAGCT |
| 12S_F | ATATACCGCCATCTTCAGCA |
| 12S_R | CTAAATCCACCTTCGACCCT |
| ATP6_F | GGACTCCTGCCTCACTCATT |
| ATP6_R | AAGTGGGCTAGGGCATTTTT |
| TFAM_F | GCTAAGGGTGATTCACCGCA |
| TFAM_R | ATCCTTTCGTCCAACTTCAATC |
| POLRMT_F | AGGTCAAGCAAATAGGAGGTG |
| POLRMT_R | CAGCGAGTGGATGAAGTTGG |
| CYTB_F | CGCATGATGAAACTTCGGCT |
| CYTB_R | ATTTGGAGGATCAGGCAGGC |
| ND3_F | GGCTTCGACCCTATATCCCC |
| ND3_R | TAGGGCTCATGGTAGGGGTA |
| SHMT2_F | CAAGACTGGCCGGGAGATC |
| SHMT2_R | GGGAACACGGCAAAGTTGAT |
| TFB1M_F | TGCTTGCCGCGTATCATG |
| TFB1M_R | CGGAGGGAGACGGCAAGT |
| MRPL1_F | TTTACAGAGAATGCATCAGAGG |
| MRPL1_R | AGGCATTATTTCTGGAACAGC |
| MRPL11_F | GAGGCGTTTCCATCAACCAG |
| MRPL11_R | CACCTCTTTCCCTGTTTGCC |
| MRPL13_F | TACTAGGAGAAGGACGTACGG |
| MRPL13_R | CACAGTCACTCAGTGCATGG |
| MRPL21_F | GGAAATGAACTAGACCTTGCGT |
| MRPL21_R | AAGATCCTTTCCGAGGAGTG |
| MRPS34_F | GTGGACTACGAGACCTTGAC |
| MRPS34_R | AAAGAGGCGTCTTTGAAGGTC |

| | |
|---|---|
| MRPS23_F | GCTCCCATCCAAGACATCTG |
| MRPS23_R | TACTTCTCCACAAACCGTTGAC |
| DDX58_F | CAAGCCTTCCAGGATTATATCCG |
| DDX58_R | AGTCCAGAATAACCTGCATGGT |
| ADAR-p150_F | CTTCCAGTGCGGAGTAGCG |
| ADAR-p150_R | GTGACGGTGTCTGCTTTCCA |
| Mx1_F | GTTACCAGGACTACGAGATTGAG |
| Mx1_R | GATGAGTGTCTTGATCTTATACCC |
| IFIT1_F | TACCTGGACAAGGTGGAGAA |
| IFIT1_R | GTGAGGACATGTTGGCTAGA |
| IFIT2_F | TGTGCAACCTACTGGCCTAT |
| IFIT2_R | TTGCCAGTCCAGAGGTGAAT |
| ISG15_F | GCGAACTCATCTTTGCCAGTA |
| ISG15_R | CCAGCATCTTCACCGTCAG |

**Supplementary figure 1. Expression of mitochondrial genes under hypoxia is not affected in 143B parental cells.** a) RNA expression of mitochondrial encoded genes (*12S, ND3, ATP6*) or nuclear encoded genes (*POLRMT, TFAM, TFB1M*) involved in mitochondrial function was evaluated by qPCR in 143B parental and Rho Zero cells cultured in normoxia or 0.1% hypoxia (n=3). Number of replicates indicate biological replicates and data is shown as mean±SEM. * p<0.05, ** p<0.01, *** p<0.001

**Supplementary figure 2. Hypoxia has a major role over the effect of different mitochondrial-targeting drugs on *IFNβ* promoter activation.** *IFNβ* promoter stimulation was evaluated in MCF7 cells cultured in normoxia or 0.1% hypoxia for 48h and treated with different drugs targeting different mitochondrial processes (n=3). Number of replicates indicate biological replicates and data is shown as mean±SEM. * p<0.05, ** p<0.01, *** p<0.001

**Supplementary Figure 1**

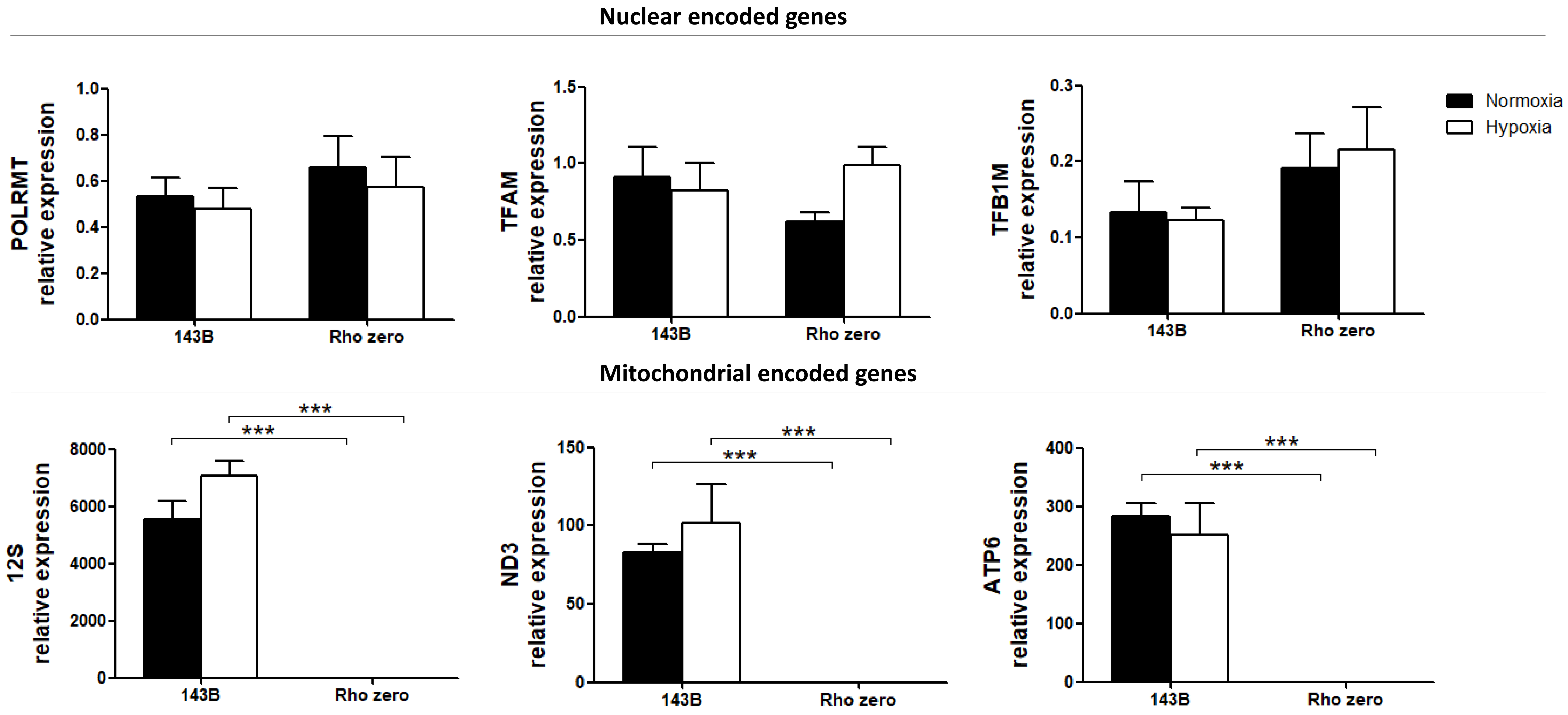

**Supplementary Figure 2**

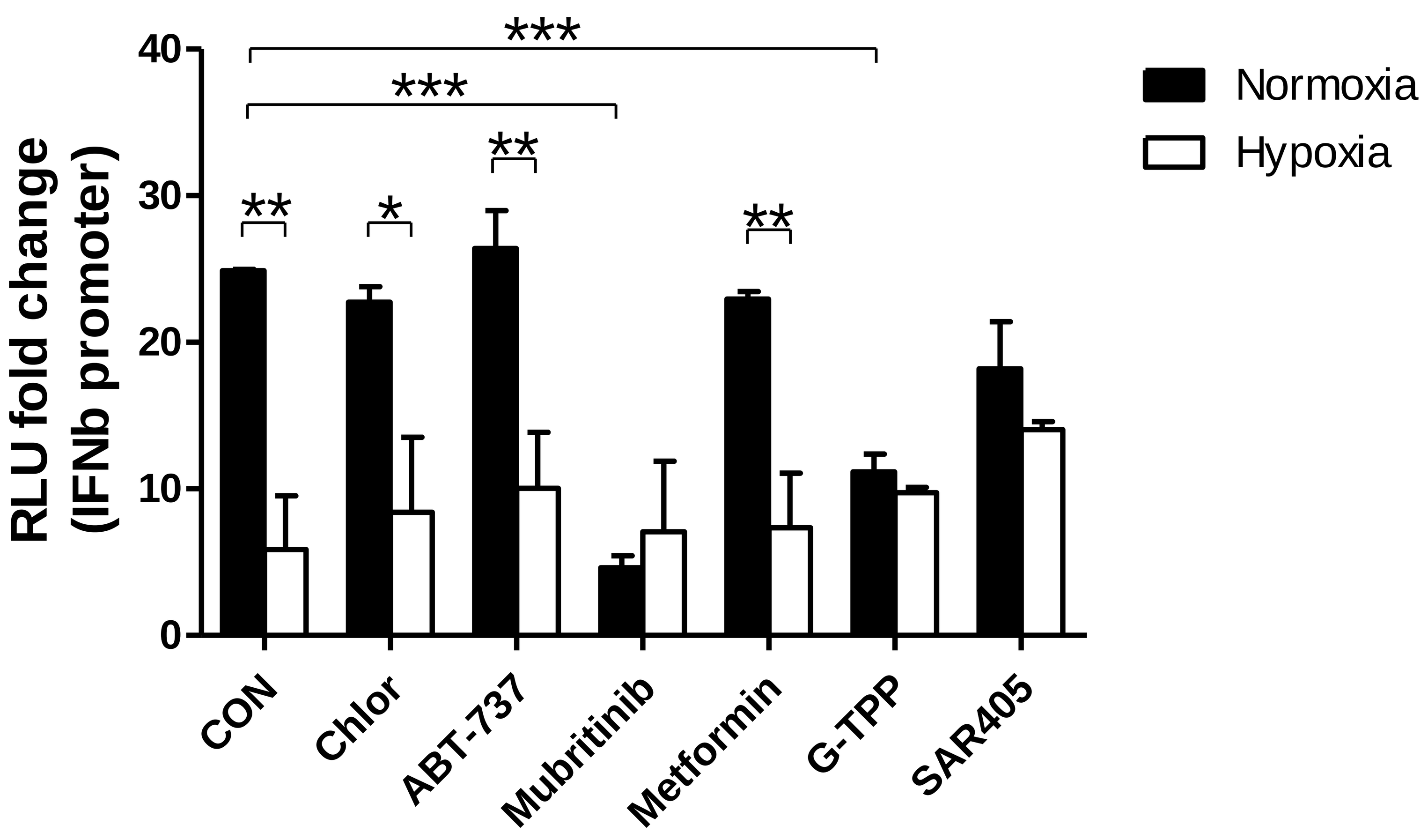
